# Mycotools: An Automated and Scalable Platform for Comparative Genomics

**DOI:** 10.1101/2023.09.08.556886

**Authors:** Zachary Konkel, Jason C. Slot

## Abstract

Comparative genomics comprises analyses that investigate the genetic basis of organismal biology and ecology, which have also been applied to high throughput trait screening for applied purposes. The number of fungal genomes deposited in publicly available databases are currently in exponential growth. Due to the limited cutting-edge software availability and size or efficiency constraints of web-based analyses, comparative genomics research is often conducted on local computing environments. There is thus a need for an efficient standardized framework for locally assimilating, curating, and interfacing with genomic data. We present Mycotools as a comparative genomics database software suite that automatically curates, updates, and standardizes local comparative genomics. Mycotools incorporates novel analysis pipelines that are built on a suite of modules that streamline routine-to-complex comparative genomic tasks. The Mycotools software suite serves as a foundation for accessible and reproducible large-scale comparative genomics on local compute systems.

## INTRODUCTION

Comparative genomic analyses are widely applied in healthcare, agriculture, and ecological studies. In healthcare, phylogenomic analysis has allowed the monitoring of strain evolution for the public health response to SARS-CoV-2 (*1–3*). At a broad taxonomic scope, phylogenomics has radically transformed the understanding of organismal relationships across life (*4*). In agricultural research, comparative genomics has unveiled population-specific virulence factors in pathogens, while genome-wide association studies have generated gene targets for breeding pest resistance (*5–7*).

The rapid accumulation of publicly available genomes, made possible by advances in whole-genome sequencing, has expanded the potential scope and resolution of comparative genomics (Figure 2.1). Web-hosted public genome resources, such as GenBank at the National Center for Biotechnology Information (NCBI) and MycoCosm at Joint Genome Institute (JGI), are central repositories for thousands of fungal genomes (*8*). GenBank’s fungal genome database appears to be in exponential growth since 2000, currently growing at a rate faster than one genome per day (Figure 1). These web databases incorporate important analysis tools, such as RefSeq *BLAST* (Madden, 2013) and the MycoCosm phylogeny (*8*) However, genomic methods have rapidly evolved in parallel with genome availability, making it impractical for web databases to implement most modern software. Therefore, online genome data is often locally assimilated and combined with genomes generated in lab in order to take advantage of cutting edge comparative genomics software.

**Figure 1:**
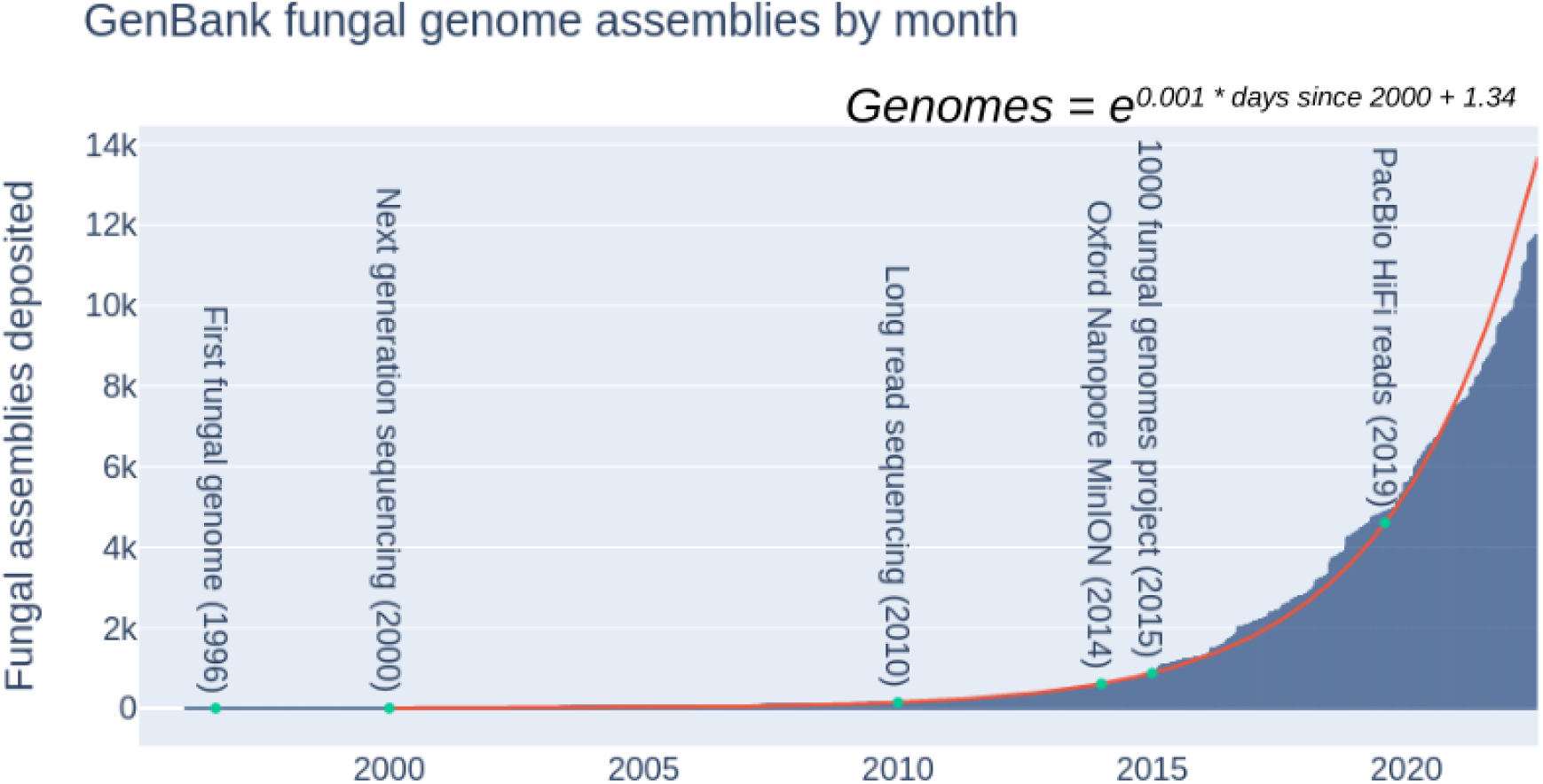
Number of publicly available fungal genomes are in exponential growth (ordinary least squares log(genomes) v. time R^2^ = 0.994).

Locally analyzing genomic data is cumbersome because standardized local database software is not broadly available. Local genome databases are built by manual curation or in-house database assimilation scripting (*9–14*) resulting in both redundancy and non-standardized file formats. Despite the large effort to create local databases, the rapid growth of available data quickly renders these databases obsolete. Acquiring available data is made more difficult by numerous legacy genome annotation file formats and discrepant formatting between different genome repositories. The complexity with assimilating genome data locally is a significant inefficiency that may lead to using rapidly outdated or incomplete datasets.

We present Mycotools, a software suite designed to increase the efficiency, accessibility, and scalability of comparative genomics by implementing a standardized comparative genomics database format and interface. Mycotools is centered around MycotoolsDB (MTDB, Figure 2), a systematically assimilated local database of publicly available genome data. MTDB automatically downloads and curates GenBank and JGI genomic data in the *mtdb* format. The *mtdb* format contains the metadata and genomic data that is required for input to the Mycotools software suite and external software. This simplifies analysis by incorporating a single input file that is also a log of the metadata associated with each analysis. Mycotools scripts and libraries are built around the *mtdb* format, which enables pipelining routine to complex comparative genomic analyses. Mycotools includes practical pipelines built from these modules, including homology search algorithms coupled with accession extraction, automated phylogenetic analysis, and synteny analysis pipelines. The Mycotools software suite is poised to serve as a foundation for a standardized comparative genomics interface.

**Figure 2:**
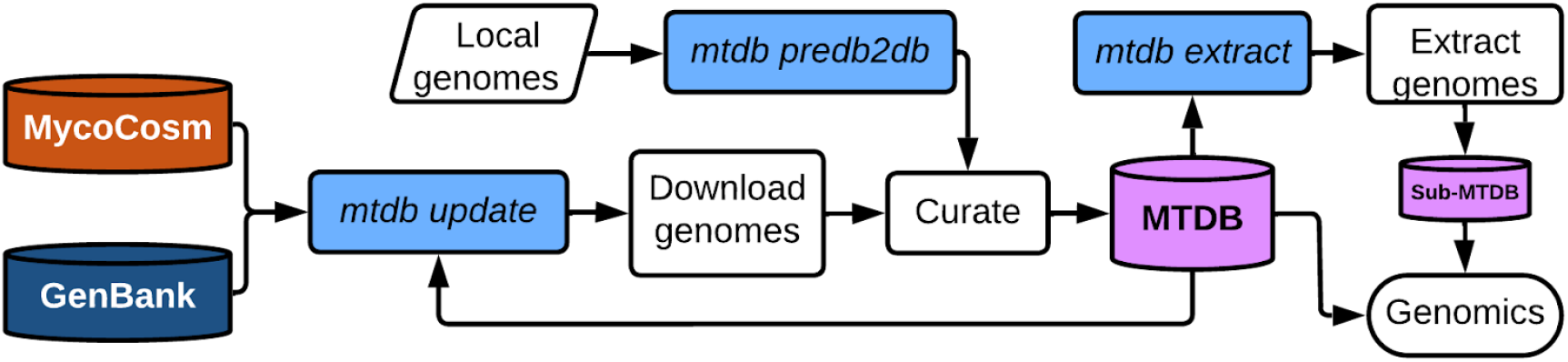
Mycotools scripts (light blue) automatically initialize and update the MycotoolsDB (MTDB), a local curated genomics database. Left to right, the command, mtdb update initializes the primary MTDB by downloading GenBank and MycoCosm genomic data; mtdb predb2db adds local genomes to the curated MTDB; mtdb update updates the database with publicly available data; mtdb extract extracts an MTDB file that is a subset of the primary MTDB for genomic analysis. The full primary mtdb is alternatively used in downstream comparative genomics.

## DESIGN AND IMPLEMENTATION

### MycotoolsDB increases the accessibility of large-scale comparative genomics

MycotoolsDB (MTDB) is designed for ease-of-use and keeping pace with accumulating genome data. MTDB is initialized and updated by assimilating publicly available genomes from GenBank and MycoCosm with local data using the script *mtdb update* (Figure 2). This script initializes by dereplicating MycoCosm and NCBI data using the assembly accession (GenBank), submitter (GenBank), date modified (GenBank), and Portal ID (MycoCosm) fields. Dereplication references an existing MTDB, the primary MycoCosm genome table and/or NCBI primary table (prokaryotes.txt/eukaryotes.txt). NCBI genomes that contain “Joint Genome Institute” or “JGI” in the submitter field are removed as redundant overlap with MycoCosm. Once data is dereplicated, Mycotools downloads the gene coordinates *gff* and masked assembly *fasta*. Each genome is systematically assigned a readable, informative “ome” code comprised of the first three letters of the genus, the first three letters of the species, and a unique accession number for that codename (e.g., the first added *Amanita muscaria* genome is denoted *amamus1*). Version updates append a version number to the code (e.g., *amamus1* to *amamus1*.*1* following the first update). Local genomes are added to the primary MTDB by submitting a genome-delimited spreadsheet of metadata, assembly paths, and gene coordinate *gff* paths, which are then curated into the same format as downloaded genomes.

Mycotools increases the accessibility and software compatibility of comparative genome analyses by systematically and uniformly curating the general features format (*gff*) annotation files and assembly *fasta*. The *gff* format needs curation because it is slow to assemble the *gff* hierarchical structure (*15*) and the format has been through three primary versions, with discrepant formatting that breaks software compatibility. Mycotools expedites *gff* parsing by curating and applying MTDB accessions to each entry, which directly links related entries without requiring hierarchical assimilation. For example, GFF data by default has to be iteratively parsed to tie CDS entries to their parent genes by first linking each CDS to their parent RNA entry. Mycotools curation additionally improves software compatibility by updating legacy *gff* versions to *gff3*, and establishing discrete, uniform guidelines for formatting. Legacy *gff* versions are brought to *gff3* by curating introns into exons and translating start and stop codon coordinates into gene entries. RNAs, exons, genes, pseudogenes, and CDSs are other acceptable *gff* entry types. The original assembly contig accessions are retained and prepended with the genome’s ome code. Full formatting requirements are detailed in the Mycotools usage guide (github.com/xonq/mycotools).

### MycotoolsDB standardizes interfacing with large-scale comparative genomics

MycotoolsDB (MTDB) genome database formatting standardizes and streamlines comparative genomics by providing a file format that can scale with the exponential growth of genomic data. MTDB files serve as the sole input that references genome data to each analysis, which increases efficiency and swift analysis scalability by extracting rows of interest for subset analyses. This format is flexible for future updates, can operate as uniform standalone reference databases, and is an easily-disseminated log of genome metadata for data reporting. MTDB files are scalable tab delimited reference files with each row containing a unique genome, its taxonomic metadata, and assembly and annotation file paths (Table 1). Mycotools analysis scripts input the primary MTDB by default, though subsets of the primary MTDB can be generated by extracting rows (genomes) of interest for an analysis. For example, a BLAST or profile model search can be executed against the primary MTDB via *db2search*, or an analysis can be restricted to a particular lineage by extracting an MTDB file of a taxonomic lineage using *mtdb extract* and inputting the extracted MTDB file to *db2search*. If external software do not have built-in compatibility for the MTDB format, reference MTDB files can still be used as a database because Mycotools curates the data and *db2files* copies requested genome data referenced in MTDB files. Due to the simplicity of the format, MTDB files are also amenable to standard shell scripting.

**Table 1:**
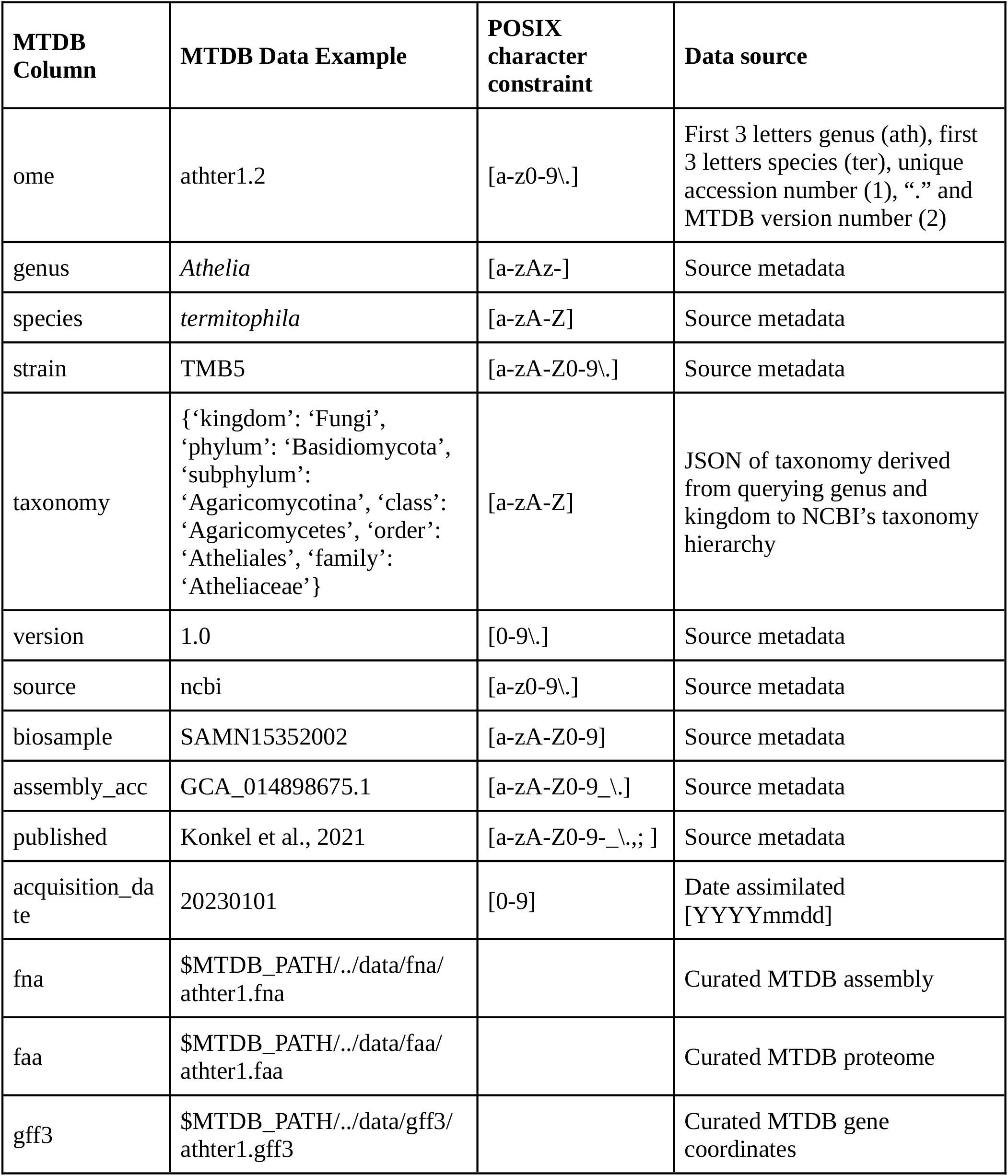
Mycotools database (MTDB) file format standard.

### Mycotools pipelining modules streamline comparative genomic analyses

Mycotools modules form the foundation for complex comparative genomics pipelines. Table 2 delineates Mycotools modules, which manipulate the *mtdb* Python object to perform routine tasks that serve as the foundation for pipelining complex analyses. For example, accession acquisition in desired formats is streamlined through the *acc2fa/acc2gff/acc2gbk/acc2locus* scripts (Table 2), while synteny diagram generation is implemented in *gff2svg*, and automated phylogenetic reconstruction that includes alignment, trimming, and tree building modules, is built into *fa2tree. fa2clus* is an example pipeline built from Mycotools modules and external dependencies, which iteratively clusters sequences based on sequence similarity until a cluster within a minimum and maximum number of sequences is obtained. *fa2clus* is additionally a module for the Cluster Reconstruction and Phylogenetic Analysis (*CRAP*) pipeline (*16*), which analyzes the evolution of a input locus by integrating locus and gene accession acquisition, homolog identification, sequence similarity clustering, phylogenetic reconstruction, and synteny analysis (Figure 3). Mycotools also implements a novel maximum-likelihood microsynteny phylogeny pipeline, *db2microsyntree*, which reconstructs a tree data structure that recapitulates divergence in gene order (*17*). When external software is used, the *db2files* module symbolically links MTDB genome data for seamless file acquisition (Figure 3).

**Table 2:**
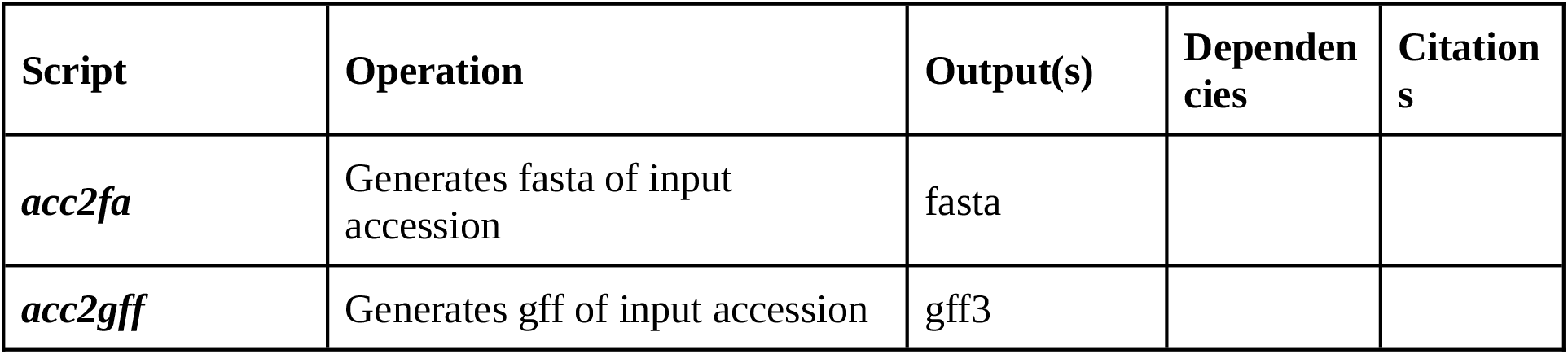
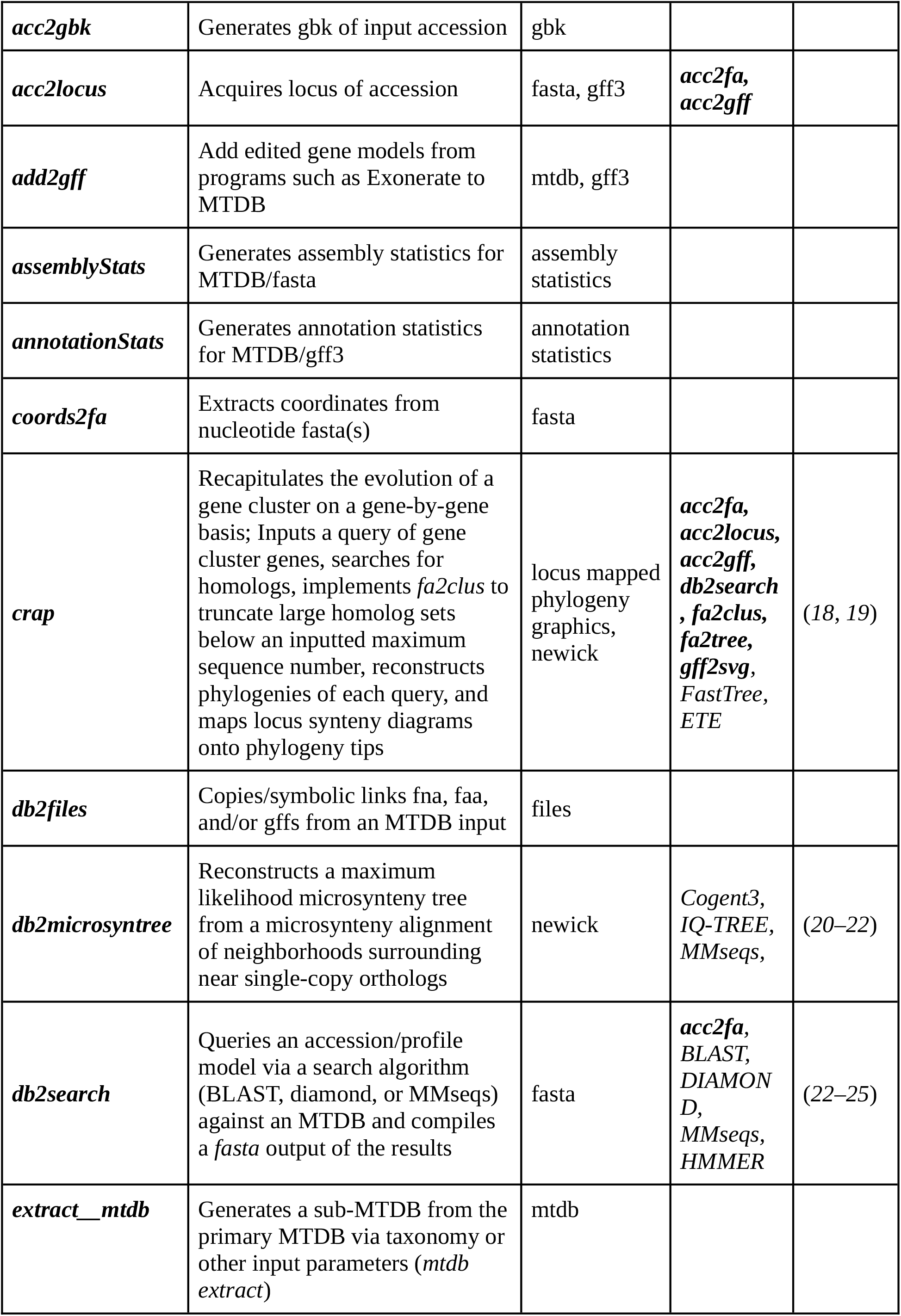
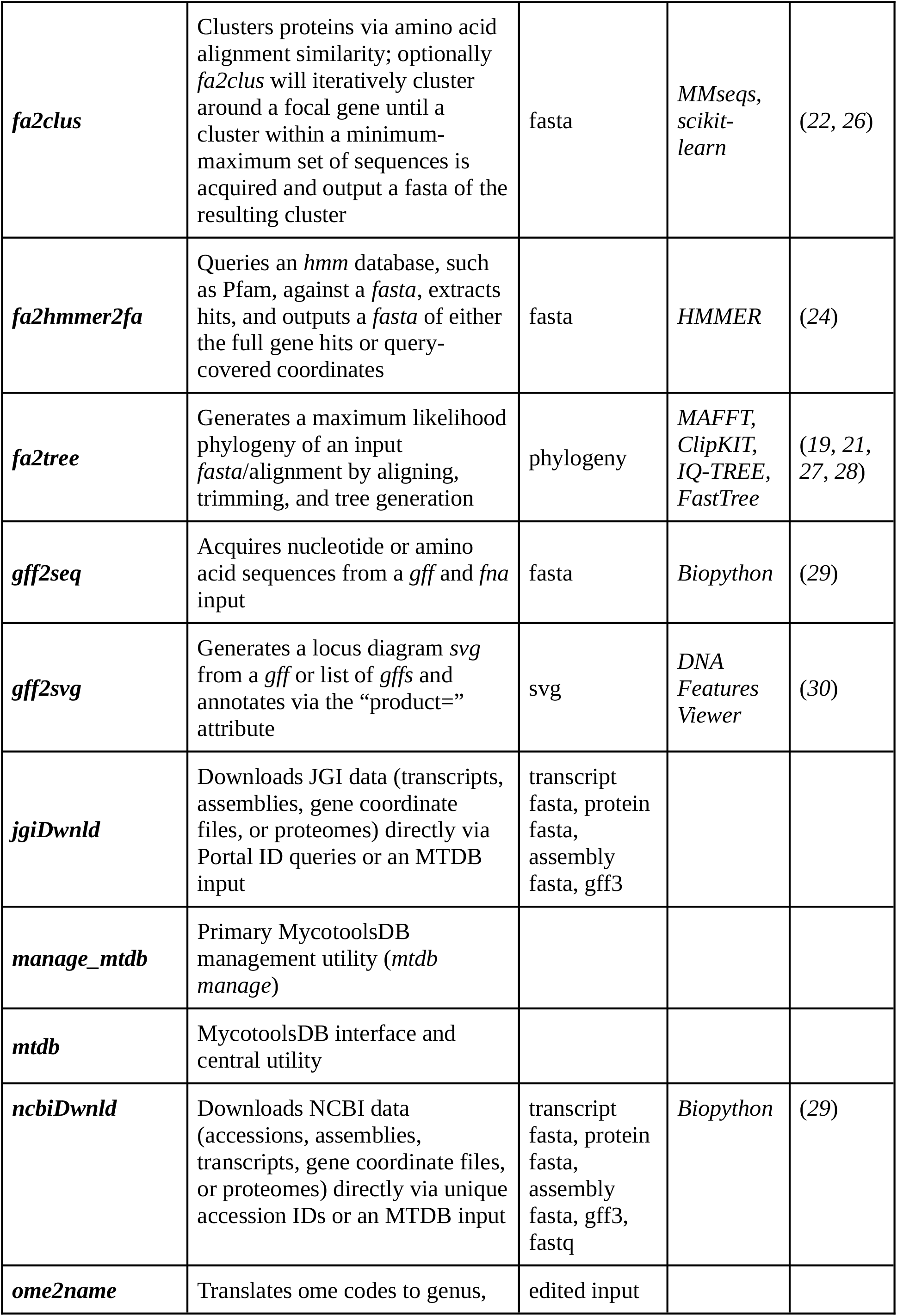
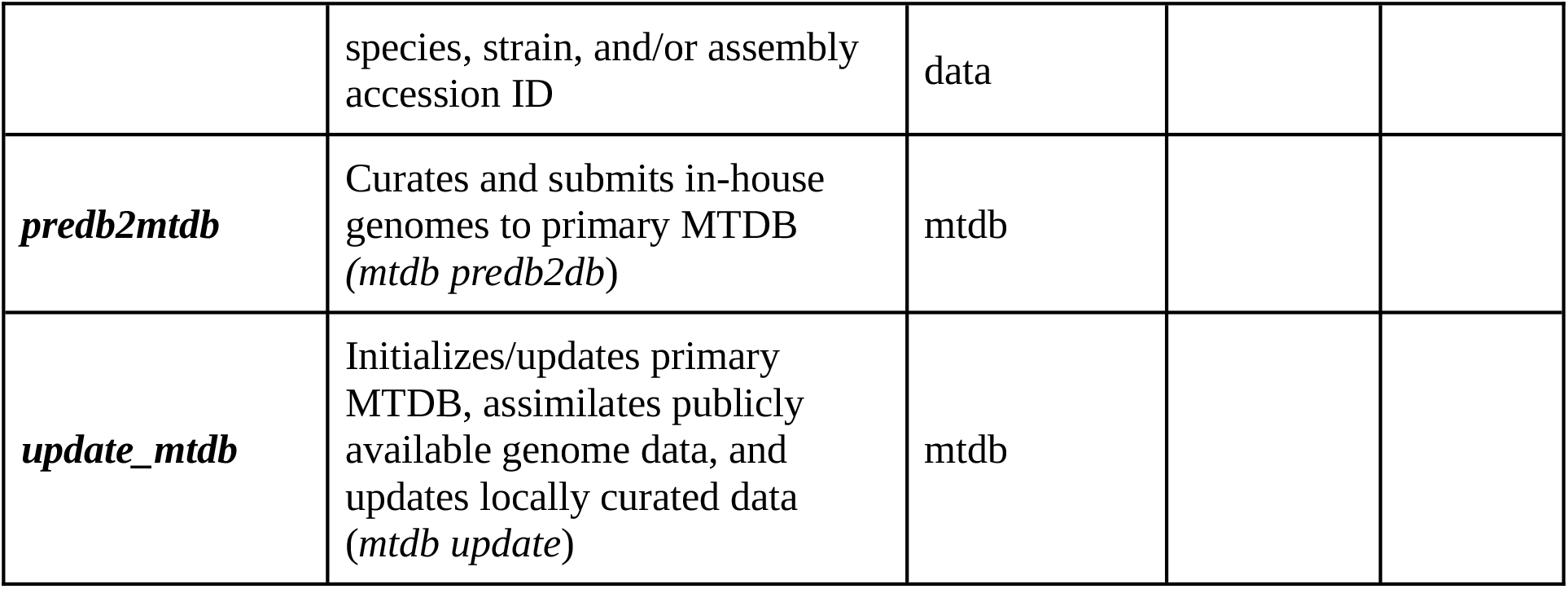
Mycotools routine analysis and pipelining modules.

**Figure 3:**
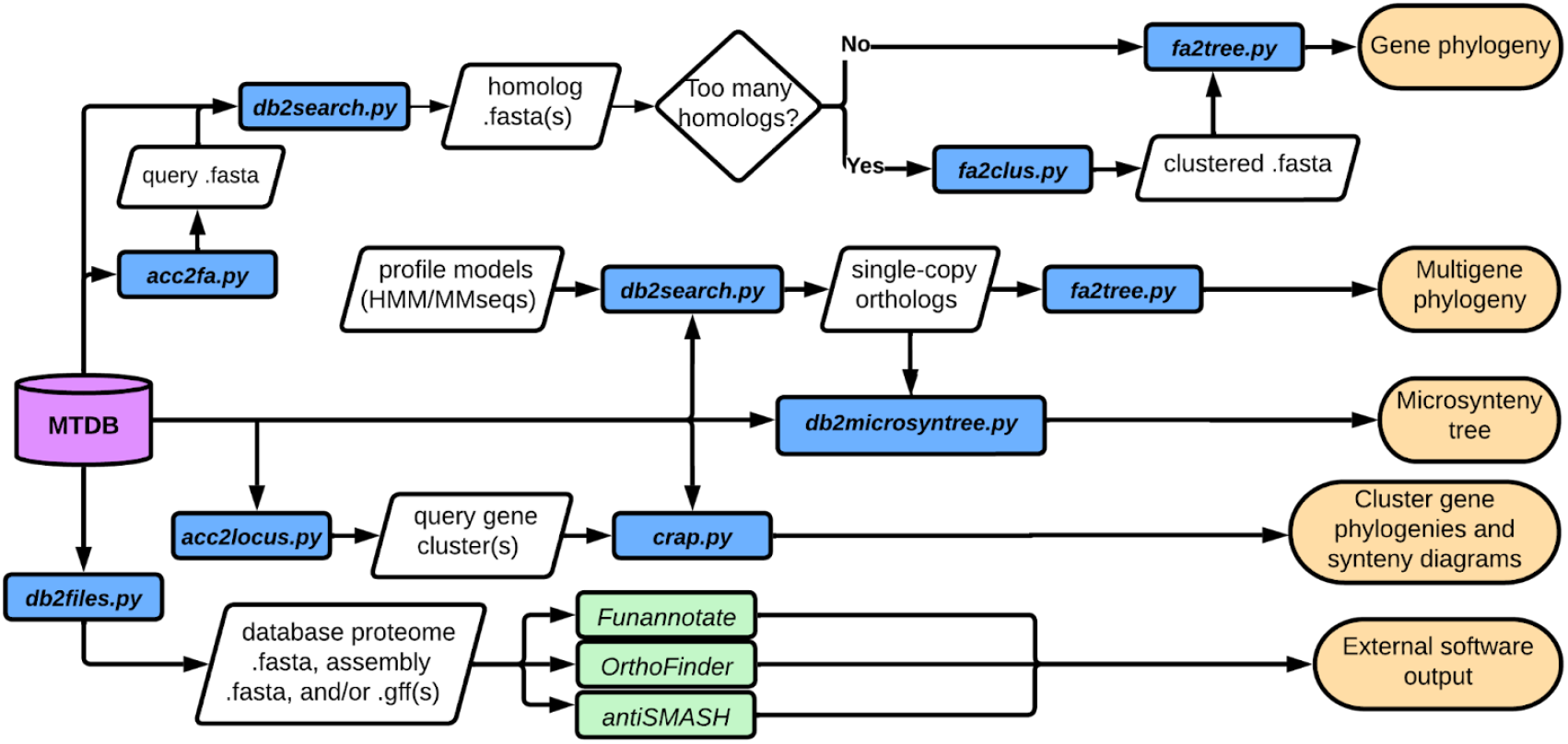
Example Mycotools scripting pipelines with Mycotools scripts highlighted in blue and pipeline output in peach. The MycotoolsDB (MTDB) is depicted in violet and represents the primary MTDB or extracted sub-MTDBs. External software are green. From top to bottom: the first pipeline illustrates building gene phylogenies, including systematically truncating large sets of homologs via sequence similarity clustering (fa2clus); multigene phylogenetic analysis is used for constructing species trees or a maximum-likelihood microsynteny tree from single-copy orthologs identified using hidden markov model searches; the Cluster Reconstruction and Phylogenetic Analysis Pipeline (CRAP) combines synteny analysis with multigene phylogenetic reconstruction, and is built on multiple Mycotools modules (Table 2); and MTDB integrates with external software by acquiring input genomic data via db2files.

### Testing Mycotools throughput using a dataset of a fungal subphylum

To demonstrate the standardization and efficiency of Mycotools-facilitated comparative genomics, we used a test MTDB of the fungal subphylum, Ustilaginomycotina (Basidiomycota). We used the Mycotools software suite to acquire genome assembly statistics, annotation statistics, reconstruct phylogenies and synteny diagrams of the nitrate assimilation gene cluster, identify single-copy orthologs, reconstruct a 14-gene species phylogeny, and generate a microsynteny tree that recapitulates gene order divergence between species. All computation was conducted using 12 Quad Core Intel Xeon 6148 Skylake processors and used 1.43 GB RAM.

## RESULTS

Mycotools automatically curates and assimilates public genomic data into a local database. The primary Mycotools-formatted database (MTDB) of all publicly-available fungi contains 3,709 fungal genomes (2023, April 11) and the prokaryote MTDB contains 267,857 (2022, September 22). Data assimilation is primarily limited by the query rate of GenBank (600 queries per minute) and MycoCosm (1 query per minute).

To demonstrate the efficiency of Mycotools for routine-to-complex genomic analyses, we performed test analyses (Figure 3) referencing 42 publicly available Ustilaginomycotina (Basidiomycota, Fungi) genomes. Each analysis is easily executed with a single command. The local assimilation of the reference dataset MycotoolsDB was completed in 83 minutes (m) using *mtdb update*. Annotation statistics for the entire database were obtained in 3 seconds (s) using *annotationStats*, while assembly statistics were gathered in 5s using *assemblyStats*. Gene phylogenies and synteny diagrams of Ustilaginomycotina nitrate assimilation gene cluster reconstructed using *crap* took 2m 23s. For generating a multigene species phylogeny, we identified 14 single-copy orthologs in all genomes in 31s using *db2search*. Then a robust multigene partition phylogeny generated using *fa2tree* with 1000 ultrafast IQ-TREE bootstrap replicates took 924m 5s. Finally, a maximum-likelihood microsynteny tree representing gene order divergence between species reconstructed using *db2microsyntree* took 2m 33s.

Beta versions of Mycotools have been implemented to facilitate novel biological discovery using large-scale local comparative genomic analysis. The Mycotools *CRAP* pipeline is a start-to-finish phylogenetic and synteny analysis pipeline (*16*)that has been used to identify horizontal transfer of the neuroactive ergot alkaloid biosynthetic gene cluster across multiple taxonomic classes (*31*). A separate analysis identified a prospective mechanism of fungal horizontal transfer by identifying transposases flanked with gene clusters using MTDB to enable large scale gene cluster identification and homology searching across a database of 1,649 publicly available fungal genomes (*5*). Mycotools also facilitated the identification of lineage-specific duplications of the industrially-important flavor genes of shiitake mushrooms by pipelining an iterative phylogenetic reconstruction analysis using 451 genomes of the mushroom-forming order, Agaricales (Basidiomycota, Fungi) (*32*). In addition to these analyses, Mycotools is directly integrated with the gene cluster detection algorithm, *CLOCI* as the source of data input for a pilot analysis of 2,247 fungal genomes (*33*). Mycotools has also been used to expedite data acquisition, genome annotation, and phylogenetic analysis of genes for novel genomes (*34–36*).

## AVAILABILITY AND FUTURE DIRECTIONS

Mycotools is freely available under a BSD 3-clause public license at github.com/xonq/mycotools. The MTDB design is taxonomy-agnostic, and is poised to suit all taxonomic domains in the future. Future iterations will improve efficiency and scalability by vectorizing serial computing functions and converting to compiled languages, such as Rust.

## ACKNOWLEDGEMENTS

We would like to acknowledge the Ohio Supercomputer Center for offering cutting-edge computing resources and Emile Gluck-Thaler, Kelsey Scott, Guillermo Valero David, Isabel Emmanuel, Lauren Slattery, Nicolle Omiotek, and Hannah Toth for implementing and testing the alpha Mycotools software suite.

